# ZEAMAP, a comprehensive database adapted to the maize multi-omics era

**DOI:** 10.1101/2020.01.04.894626

**Authors:** Songtao Gui, Linfeng Yang, Jianbo Li, Jingyun Luo, Xiaokai Xu, Jianyu Yuan, Lu Chen, Wenqiang Li, Xin Yang, Shenshen Wu, Shuyan Li, Yuebin Wang, Yabing Zhu, Qiang Gao, Ning Yang, Jianbing Yan

## Abstract

As one of the most extensively cultivated crops, maize (*Zea mays* L.) has been extensively studied by researchers and breeders for over a century. With advances in high-throughput detection of various omics data, a wealth of multi-dimensional and multi-omics information has been accumulated for maize and its wild relative, teosinte. Integration of this information has the potential to accelerate genetic research and generate improvements in maize agronomic traits. To this end, we constructed ZEAMAP (http://www.zeamap.com), a comprehensive database incorporating multiple reference genomes, annotations, comparative genomics, transcriptomes, open chromatin regions, chromatin interactions, high-quality genetic variants, phenotypes, metabolomics, genetic maps, genetic mapping loci, population structures and domestication selection signals between teosinte and maize. ZEAMAP is user-friendly, with the ability to interactively integrate, visualize and cross-reference multiple different omics datasets.

## Introduction

Maize (*Zea mays* L.) is one of the most important crops for food, feed and fuel, and is also a model species for genetic and genomic research. As the cost of sequencing has been decreased and new omics technologies have arisen, there has been explosive growth in the amount of biological information available for maize. The maize B73 reference genome has recently been updated (1), and four more maize genome assemblies have been released during the last two years (2-5). The previous two-dimensional genome has recently been resolved in three dimensions with the mapping of open chromatin and the identification of chromatin interactions based on ChiA-PET and Hi-C technologies (6,7). Omics data, including deep DNA resequencing, transcriptome and metabolome, have been accumulated at the population scale (8-15). There are many different applications for these new data sets, including gene cloning and the study of regulatory networks. These new and comprehensive data sets provide valuable resources for maize research, and have the potential to completely revolutionize breeding (16).

Comprehensive databases are needed to store, maintain and analyze the multi-omics data which is now available for maize. Several maize genomics and functional genomics databases have been developed, including the Maize Genetics and Genomics Database (MaizeGDB) (https://www.maizegdb.org/), which collects maize reference sequences, stocks, phenotypic and genotypic data and also provides useful tools for maize data mining (17,18). Panzea (https://www.panzea.org/) collects genotypic and phenotypic information for several maize populations (19). while MaizeNet (http://www.inetbio.org/maizenet/) provides a genome-scale co-functional network of maize genes (20). Other generic databases such as Genbank (https://www.ncbi.nlm.nih.gov/genbank/), Gramene (http://www.gramene.org/) and ePlant (http://bar.utoronto.ca/eplant_maize/) also collect maize omics data. Despite being very useful, these databases are designed either to collect general maize genomic and genetic information or to focus on one specific omics area. To make the best use of the multi-omics information for maize research and breeding, researchers currently need to either systematically integrate omics data generated from different sources (21), or use multi-omics data which were all generated from the same panel.

MODEM (http://modem.hzau.edu.cn/) is the first attempt to integrate multi-omics datasets, including various types of genetic variants, expression data and metabolomic data (22). Despite the importance of wild relatives in understanding the domestication of modern crops, no existing maize multi-omics databases incorporate teosinte (23). To fill this gap, we have developed ZEAMAP (http://www.zeamap.com/), a multi-omics database for maize research and breeding, which integrates omics data generated from 507 elite inbred lines (in an association mapping panel, AMP) (24) and 183 teosinte accessions. ZEAMAP includes genome assemblies and annotations of four inbred lines, B73 (1), Mo17 (3), SK (4) and HuangZaoSi (HZS) (5), and a teosinte accession (*Zea mays* ssp. *mexicana*) (25), expression patterns of tissues from different development stage of same inbred line (4,9) and same tissue of different samples within the AMP (10), three dimensional chromatin interactions and open chromatins of B73 (6), genetic variations including single-nucleotide polymorphisms (SNPs), small insertions and deletions (InDels) and large structure variations (SVs) generated from the deep sequencing of the AMP and the comparison among reference genome assemblies, the phenotypes and metabolome of the AMP and the related loci mapped by genome-wide association studies (GWAS), expression quantitative trait locus (eQTL) and linkage analysis, the population structure and pedigrees of each germplasm and the selective signals between different teosinte subspecies and maize. ZEAMAP generated comprehensive functional annotations for the annotated gene models in each assembly, and provided useful tools for users to search, analyze and visualize all these different omics data.

## Materials and methods

### Data collection

The pedigree information of elite inbred lines in the maize association mapping panel was collected from (24). The B73 reference genome assembly (AGPv4) and annotation files (Version 4.43) were downloaded from Gramene (https://www.maizegdb.org/genome/genome_assembly/Zm-B73-REFERENCE-GRAMENE-4.0), while the Mo17 reference genome (version 1.0) and annotations (version 1.0) were downloaded from MaizeGDB (https://ftp.maizegdb.org/MaizeGDB/FTP/Zm-Mo17-REFERENCE-CAU-1.0/). The SK reference genome (version 1.0), annotations (version 1.0) and RNA-seq data of nine SK tissues were collected from (4). The HZS genome assembly, annotations and RNA-sequencing data were retrieved from Genome Sequence Archive in Beijing Institute of Genomics (BIG) Data Center (http://bigd.big.ac.cn/gsa) with project ID PRJCA001247. The genome assembly and annotations of *Zea mays* ssp. *mexicana* were from (25). RNA-sequencing of different B73 tissues were from (9), and RNA-seq data of developing maize kernels from 368 AMP inbred lines were from (10). The chromatin interaction and histone modification data were from (6), the chromatin accessibility data were from (7), and the DNA methylation data of the AMP were from (26). Phenotypes of the AMP, including agronomic traits, kernel amino acid contents, kernel lipid contents and metabolomic data were collected from previously reported studies (11,27,28). Linkage maps and QTL mapping results were collected from (29).

The SNPs and InDels were identified and genotyped using DNA deep sequencing data (∼20x) of the 507 maize AMP inbred lines (NCBI Bioproject: PRJNA531553) (4) and 183 teosinte samples (unpublished). The eQTLs, the population structures and the selective signals were generated based on these populational sequencing data. A related research article with accessions and details of the deep sequencing data, the SNPs and InDels, the eQTLs, the population structures and the selective signals is being prepared.

### Functional annotation

For each genome annotation, the protein sequences of the predicted genes were compared against the InterPro database using InterProScan 5 (30) to identify functional protein domains. The proteins were further compared against the GenBank non-redundant protein (nr) database using Basic Local Alignment Search Tool (BLAST) with the options “-p blastp –e 1e-05 –b 5 –v 5 –a 4 –m 7 –F F”. The BLAST results against the nr database and the Interpro results were further analyzed by Blast2GO (31) to assign gene ontology (GO) terms. Kyoto Encyclopedia of Genes and Genomes (KEGG) annotations were performed by running BLAST against the KEGG database (version 84.0) with options “-p blastp -e 1e-05 -a 4 -m 8 -F F”. The proteins were also searched against PFAM version 32.0 (32) using HMMer 3.1b2 (33) with default parameters. To identity gene orthologs and clusters of orthologous group (COG) annotations, the proteins were mapped to eggNOG orthology database (version 4.5.1) (34) using emapper-1.0.3 (35). To add gene-product annotations, the proteins were searched against UniProt database (version 2019_04) (36) using Diamond (v0.8.22.84) (37) with the options “--evalue 1e-05 --max-target-seqs 1”, the UniProt and EggNog search results were combined to get the gene and product names using Gene2Product v1.32 (https://github.com/nextgenusfs/gene2product). Possible proteolytic enzymes were annotated by searching the proteins against the MEROPS database (version 12.0) (38) using Diamond with the options “--evalue 1e-05 --max-target-seqs 1”. The proteins were also searched against the BUSCO (version 2.0) (39) dikarya models using HMMer with default options.

### Comparative genomics

To identify synteny blocks, we first compared proteins from one genome to those from another using BLASTP with an E-value cutoff of 1e-10 and a maximum number of alignments of 5. The significant hits were then analyzed by MCScanX (40) with parameters “-k 50 -g -1 -s 5 -e 1e-10 -m 25 -w 5” to obtain synteny blocks. The whole genome alignments between two genomes in ZEAMAP were performed using minimap2 (version 2.17-r941) (41), with parameters “-c -x asm5 -B5 -O4,16 --no-long-join -r 85 -N 50 -s 65 -z 200 --mask-level 0.9 --min-occ 200 -g 2500 --score-N 2”, and the raw alignment results were filtered to get the best alignment for each contig with QUAST-LG (42).

### Annotating of genetic variations

The SNPs and InDels were annotated using the Ensembl variant effect predictor (VEP) (43) according to B73 gene annotation v4.43. The polymorphic SVs between B73 and SK, as well as their genotypes in the AMP, were retrieved from (4), and annotated according to B73 gene annotation v4.43 using SURVIVOR v1.0.6 (44). Haplotype blocks and tag SNPs were identified using Haploview (45).

### Mapping and filtering of genetic loci

To perform genome-wide association studies for the collected phenotypic traits, the SNPs were then further filtered to keep only records with a minor allele frequency (MAF) of at least 5%. A mixed linear model accounting for the population structure (Q) and familial relationship (K) was used to examine the association between the SNPs and each trait using Tassel3 (46). The *P* value of each SNP was calculated, and significance was defined with Bonferroni corrected *P* value cutoff of 1/N, where N is the total number of markers used. To prevent the interactive GWAS viewers and the tabular loci browser from operating too slowly, the volume of GWAS results was reduced by filtering out SNPs which had very low significance values (*P* value > 1e-4). The pairwise LD r^2^ values of the remaining SNPs for each trait within 500 Kb windows were calculated using PopLDdecay (47).

We kept *cis*-eQTLs alone by retaining only the SNPs within 1 Mb of each gene (48). High quality *cis*-eQTL SNPs were selected by only retaining those with a *P* value smaller than the Bonferroni corrected *P* value cutoff of 1/N. The pairwise LD r^2^ values of the remaining SNPs for each gene were calculated using PopLDdecay (47).

### Interactive visualization tools

The visualization tools for GWAS results and selection signals were developed using LocusZoom.js (https://github.com/statgen/locuszoom), a JavaScript embeddable plugin for interactively visualizing statistical genetic data, and ECharts (https://www.echartsjs.com), an open-sourced JavaScript visualization tool. The gene expression pattern viewer and the eQTL visualizer were modified from GTEx visualizations (https://github.com/broadinstitute/gtex-viz) (48). The principal component analyses (PCA) dot plot and the ancestries stacked histogram were also developed using ECharts.

### CRISPR/Cas9 single-guide RNA designing

CRISPR/Cas9 sgRNAs for each maize reference genome were designed using CRISPR-Local (49) with default options. Results were converted into gff format with in-house perl scripts to format them for JBrowse.

## Database contents and features

### Overview structures

ZEAMAP is comprised of a user account management system, a main database, a full-site search engine, and a set of analysis and visualization tools (Figure S1). The multi-omics data in ZEAMAP are categorized into six main content modules involving genomic, genetic, variation, population, evolution and epigenetic information (Figure 1). ZEAMAP construction utilized the biological community database construction toolkit Tripal (50), which combines the content management system Drupal (https://www.drupal.org) with the standard biological relational database storage backend, Chado (51). Each feature in ZEAMAP has its own page and features are linked to each other by sequence ontology relationships.

**Figure 1.**
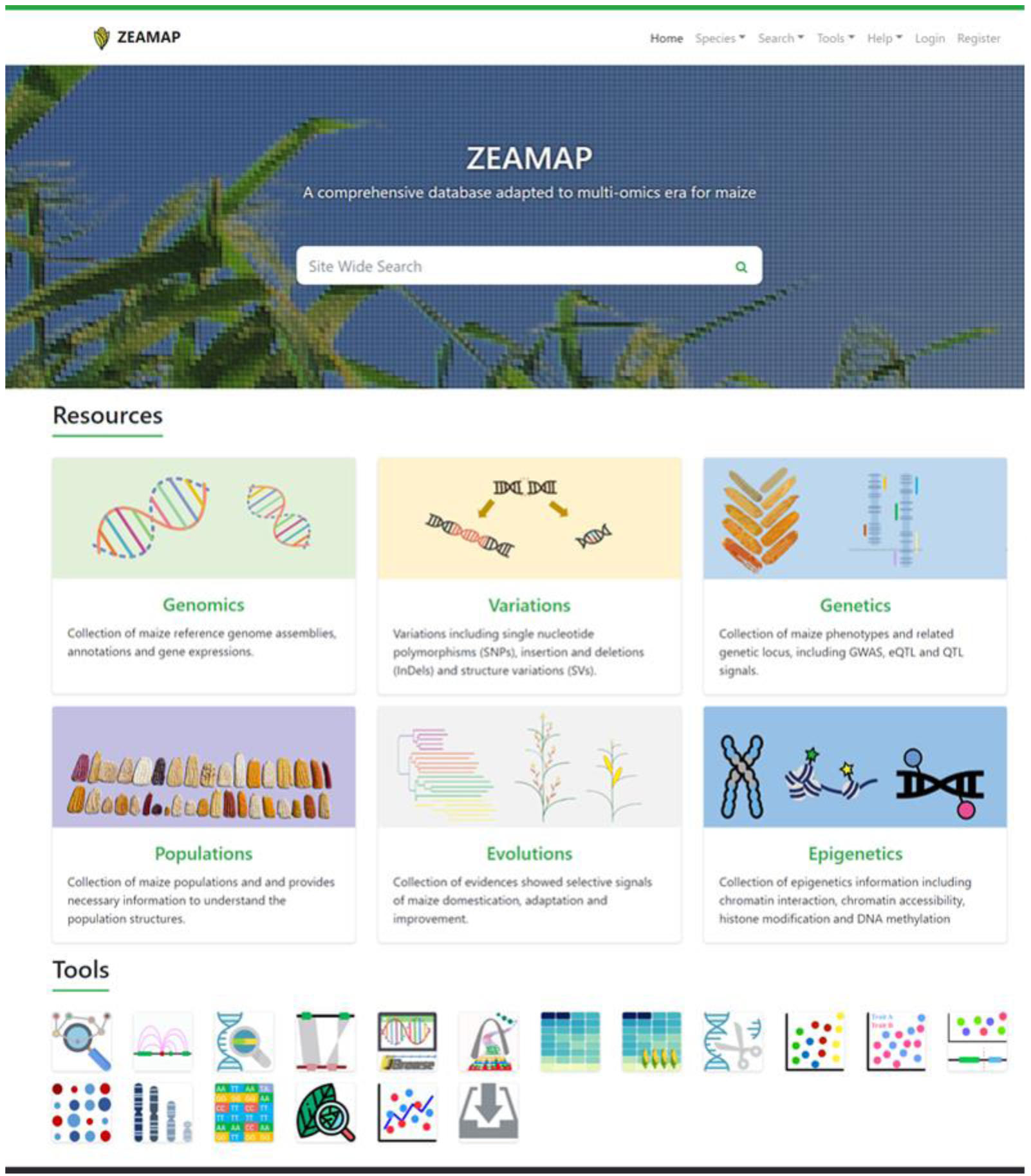
A screenshot of the ZEAMAP home page. The home page of ZEAMAP consists of a top menu bar, a site-wide search engine, access to the six biological modules and miscellaneous tools.

### The Genomics module

The Genomics module collects reference genome assemblies, gene expression profiles and comparative genomics information related to the available genomes and populations in ZEAMAP. Currently, ZEAMAP contains reference genome assemblies of four maize inbred lines (B73, SK, Mo17 and HZS) and one teosinte species (*Zea mays ssp.* mexicana). Each genome assembly has its own page which contains general information for each genome assembly and sub-menus with links to access various related information and bioinformatic analysis tools (Figure S2). The mRNA and predicted protein for gene models in each genome assembly were assigned functional annotations including gene ontologies (GO), Kyoto Encyclopedia of Genes and Genomes pathways (KEGG), clusters of orthologous groups (COG), orthology groups, known gene-product annotations, proteolytic enzymes and homologs found in InterPro, PFAM, BUSCO and NCBI nr databases. The genome features for each assembly, including genes, mRNAs, proteins and transposable elements, can be searched by their IDs or locations through Chado feature search (Figure S3A). Genes (as well as mRNAs and proteins) can also be searched by their functional annotation descriptions (Figure S3B). Each annotated genome feature has its own page with multiple sub-menus displaying summary information (resource type, accession, organism, name, identifier and others), sequences, annotations, cross references linked to the same feature in MaizeGDB or NCBI, as well as related parent and child features, orthologs and synteny blocks.

Two genome browsers, JBrowse (52) and WashU Epigenome Browser (53), were embedded to display the genome sequences, annotated genomic features and other genomic information for all the available reference genome assemblies in ZEAMAP (Figure 2A). Both genome browsers are designed to easily add tracks, search for certain information in specific regions and export data as well as figures. These two genome browsers share some common features, including genome sequences and genomic annotations. However, each one has unique information tracks (see sections below) because JBrowse performs better when dealing with large piecemeal features such as variations and the WashU Epigenome Browser is specially designed to display epigenomic tracks.

**Figure 2.**
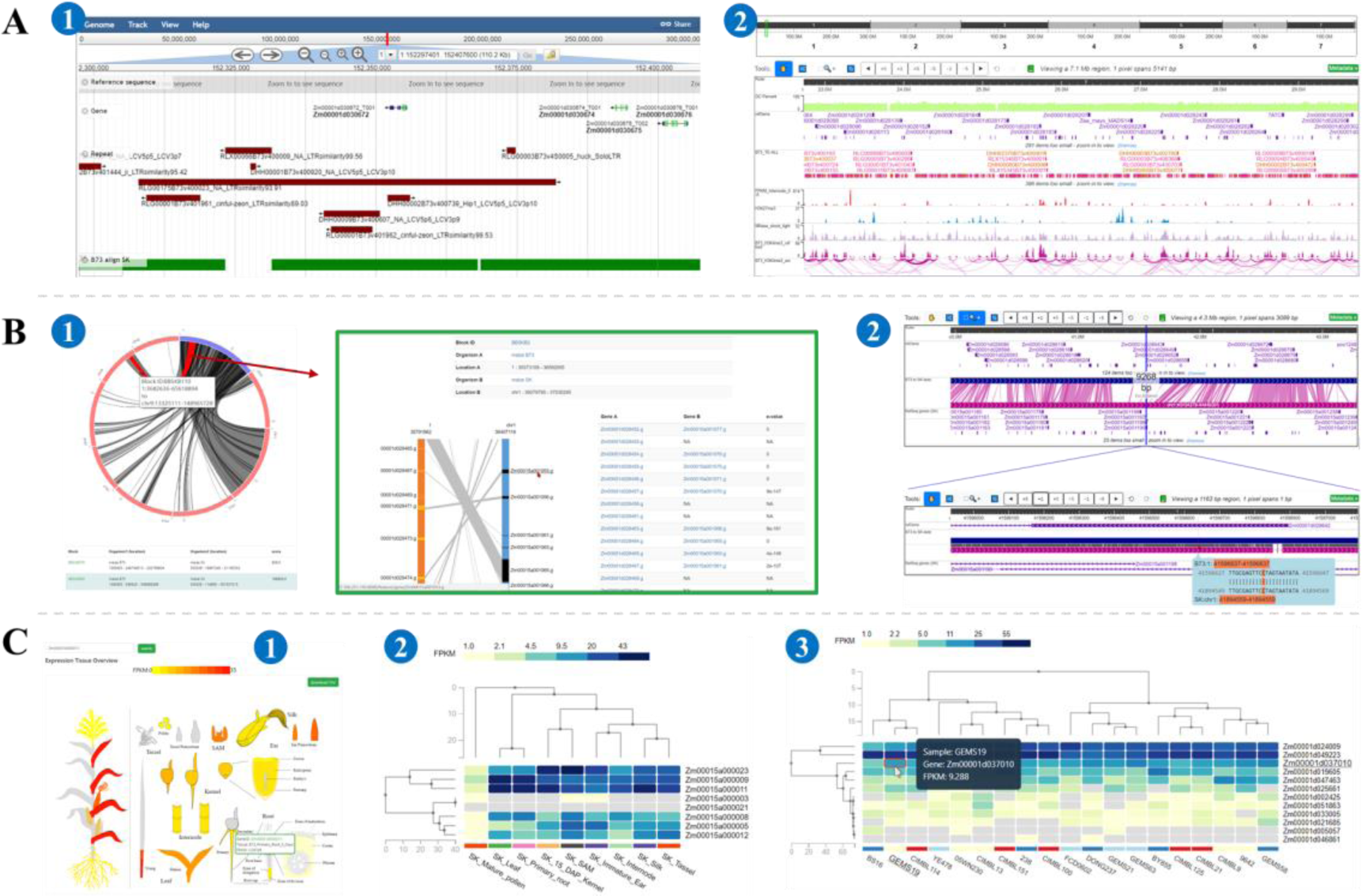
Features of the ZEAMAP genomics module. (A) Schematic of the two genome browsers embedded in ZEAMAP, JBrowse① and the WashU Epigenome Browser②. (B) Comparative genomic information in ZEAMAP, including Gene synteny blocks displayed by the synteny viewer with interactive circos plots and links to the detailed collinearity of the included genes①. Whole genome sequence alignments between two genomes are also accessible through the WashU Epigenome Browser, with Zoom-In and -Out functions and mouse-over display of the detailed alignments②. (C) Gene expression functions in ZEAMAP. The Tissue Overview function shows the expression of a gene in different tissues, with more detailed information available upon click①. ZEAMAP also has functions to cluster and display the expression patterns of several genes by tissue type② or sample③, with the gene IDs linked to pages with more detailed information.

ZEAMAP provides comparative genomic information for each pair of the available reference genome assemblies, including both synteny blocks identified from gene collinearities and whole genome alignment details. The synteny blocks are managed and displayed through the Tripal synteny viewer module (https://github.com/tripal/tripal_synview). Each synteny block has its own unique block ID and can be searched by it in both the Synteny block browser page and in the full-site search engine (Figure 2B). The detailed whole genome sequence alignments can also be accessed through the genome browser (Figure 2B).

We have collected gene expression patterns in different tissues for each maize genome assemblies, as well as expression profiles of kernels for 368 inbred lines of the AMP (10) based on B73 reference annotations. Expression patterns in different tissues for each gene can be visually displayed through heat maps after being queried in the “Tissue Overview” page (Figure 2C). ZEAMAP also enables users to browse the expression patterns of several genes among different tissues or samples and cluster the genes and tissues/samples based on the gene expression patterns (Figure 2C). Both functions provide download links to a raw expression matrix of the queries.

### The Variations module

The Variations module collects the genotypes and annotations of polymorphic variations including SNPs, InDels and SVs among the AMP in reference to the B73 reference genome, as well as a haplotype map generated from the SNP genotype matrix (See Materials and methods section for the source and the analysis used to generate the related data). The general variation information of a gene, including variation positions, allele types and annotations can be queried by their IDs or locations and displayed through tabular view (Figure S4A). The variations can also be browsed through JBrowse. Upon clicking each variant block in JBrowse tracks, the detailed information about that variant, including the annotations and the genotype of each germplasm will be shown. There is also a genotype overview for the variations in the current JBrowse display panel when the related “variant matrix” track is selected (Figure S4B). ZEAMAP also provides a function to query for the detailed genotype matrix for specified germplasms within certain regions (Figure S4C).

### The Genetics module

ZEAMAP has collected phenotypic data from the AMP, including 21 agronomic traits, 31 kernel lipid content-related traits, 19 kernel amino acid content-related traits and 184 known metabolites of maize kernels. All these phenotypes can be searched and filtered by their threshold values using the “Search Trait Evaluation” tool (Figure S5). We have identified loci significantly associated with these phenotypes using GWAS and provided a tabular data search function to find specific loci by trait names, variant IDs, chromosome regions and significant *P* values (Figure S6). Three GWAS visualization tools (“GWAS-Single-Trait”, “GWAS-Multi-Trait” and “GWAS-Locus”) were developed to better browse the GWAS results and compare the significant signals among different traits. Querying a trait and navigating to specific regions can be easily accomplished by inputting boxes or through the interactive navigational Manhattan plot (in “GWAS-Single-Trait” and “GWAS-Multi-Trait” tools). The GWAS-Single-Trait tool displays all signals associated with the selected trait as a scatter plot, with colors indicating the linkage disequilibrium (LD) r^2^ values between the user-selected reference variant and all the other variants (Figure 3A). The GWAS-Multi-Trait tool was designed to compare GWAS signals among two or more traits, with the colors indicating different traits (Figure S7), while the GWAS-Locus tool displays GWAS signals of all traits that show significant association with the query variant (Figure S8). These three tools are provided with a lightweight genome browser which indicates the gene models within the current region. Each element in the plot is also interactive and links to other related information.

**Figure 3.**
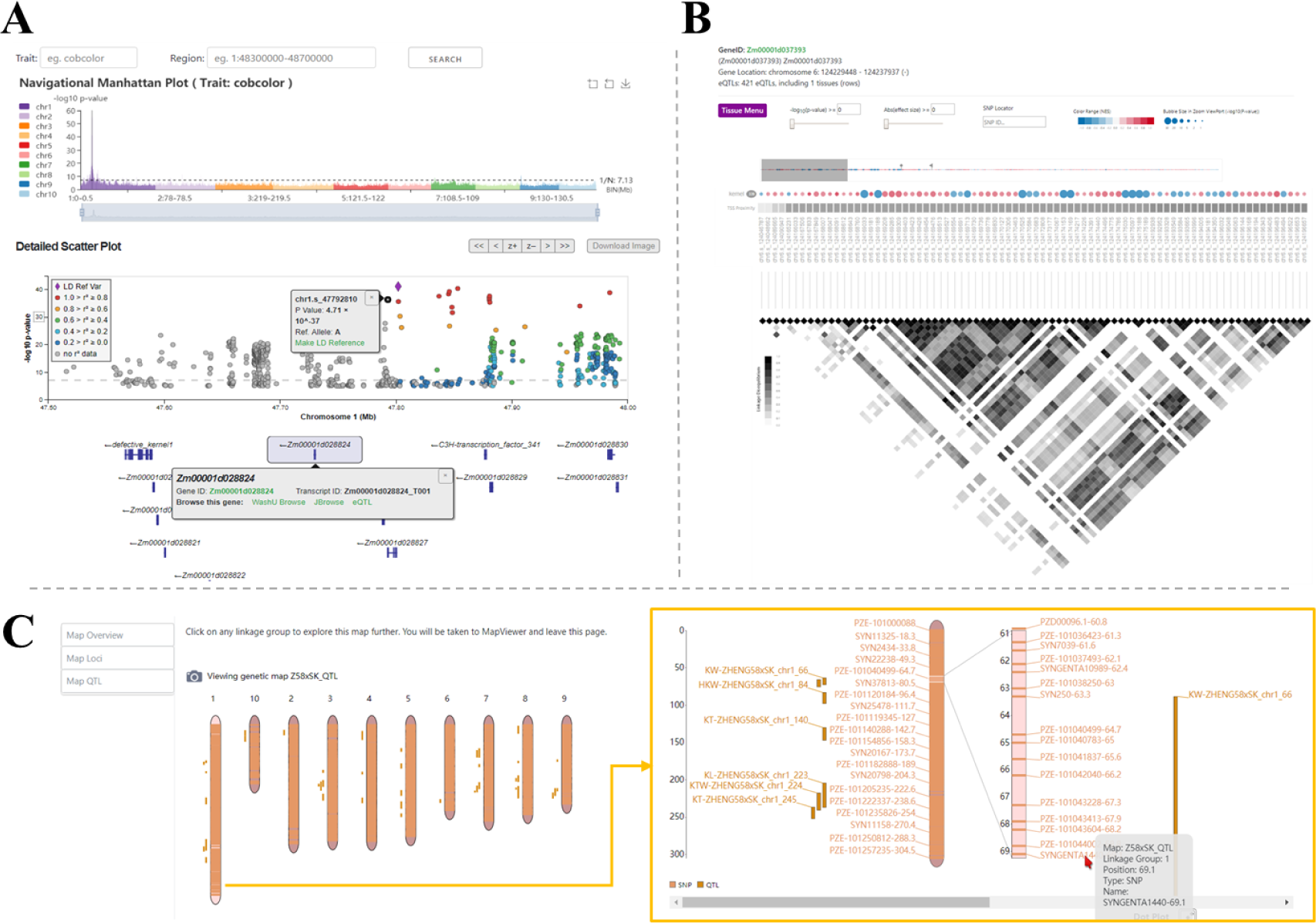
Features of the ZEAMAP genetics module. (A) Schematic of the “GWAS-Single-Trait” tool. The trait and region of interest can be queried through the top input boxes. Regions can be easily browsed through by clicking on the histogram of the interactive “Navigational Manhattan Plot” track. The “Detailed Scatter Plot” track plots the variants according to their chromosome locations and by the significance of their *P* values. The colors of each dot indicate the LD r^2^ values between that variant and the reference variant (the purple diamond dot, can be reset by selecting the “Make LD reference” link on the popup page for each variant). The bottom track shows the gene annotations in the selected region, with a popup for each gene element which links to a detailed information page, genome browsers and the eQTL visualizer for that gene. (B) Schematic of the eQTL visualization tool. The significant *cis*-eQTL site for each gene is sized by the significance of its *P* value and colored by the effect size (beta value). The heatmap indicates pairwise LD r^2^ values of the variants. (C) Schematic of the TripalMap tool in ZEAMAP. This tool displays the detailed genetic markers and mapped QTLs for each linkage group. Both the markers and the QTLs link to their own detailed information page.

Genetic variations can impact gene expression through many factors, including alterations in splicing, noncoding RNA expression and RNA stability (54). Expression quantitative trait locus (eQTL) mapping is a powerful approach to detect the possible variants which alter gene expression. In ZEAMAP, we have collected *cis*-eQTL signals with gene expression patterns in maize kernels based on B73 annonation, and provided a tabular tool to search and filter eQTL signals by gene IDs, gene locations, distances from transcription start site (TSS), effect sizes, and significance values (Figure S9). A visualization tool was also developed to browse all *cis*-eQTLs affecting the selected gene, with significance values, effect size and pairwise LD information displayed interactively (Figure 3B).

ZEAMAP has currently collected 12 published genetic maps constructed from different artificial maize segregating populations using genotypes generated from the Illumina MaizeSNP50 BeadChip (Illumina Inc., San Diego, CA, USA), as well as 813 quantitative trait loci (QTLs) identified from 15 plant architecture-related traits (29). The genetic markers can be searched and filtered by their IDs, genomic locations and genetic linkage group (Figure S10). QTLs can be searched by traits and QTL labels, resulting in detailed records of the genetic markers located in or adjacent to that QTL. By employing the TripalMap extension module (https://github.com/ksbuble/TripalMap), the linkage maps, including all related markers and QTLs, can be visualized and compared with another map interactively (Figure 3C).

### The Populations module

It is often useful to dissect the genetic diversity, population structure and pedigrees of maize lines for both evolutionary studies and molecular breeding. ZEAMAP provides interactive information about the population structures assessed by principal component analysis (PCA) and ancestries inferred from an unsupervised clustering analysis for the whole *Zea* population and each sub-population in the database (Figure 4A). We have also added a table that lists the origins or pedigree information for each inbred line of maize AMP (Figure 4B).

**Figure 4.**
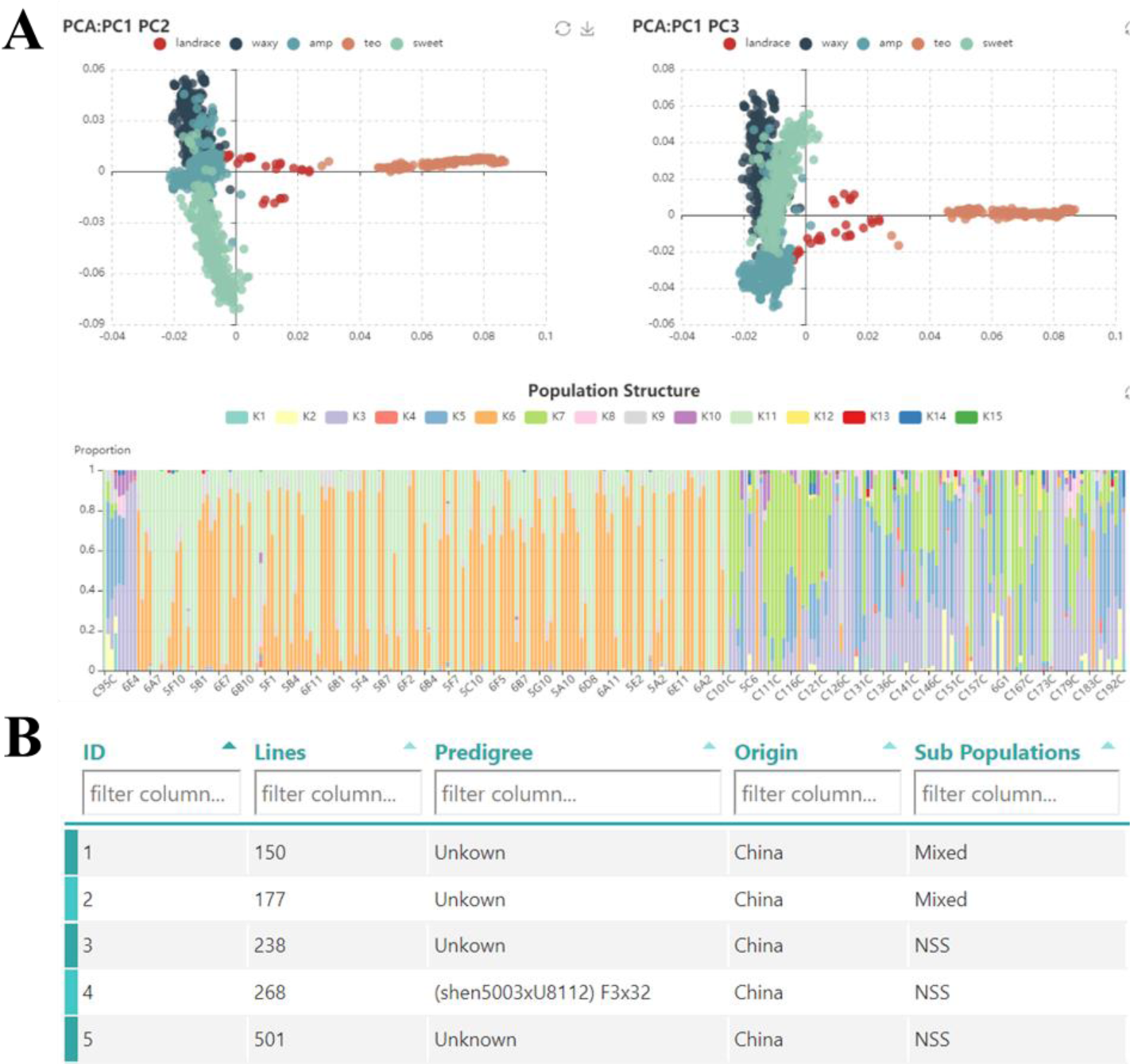
Features of the ZEAMAP Populations module. (A) Interactive PCA diagram (top two dot plots) and structure diagram (stacked bar plot). Each diagram is zoomable and shows detailed information, including germplasm names and PCA/structure values when an element is moused over. (B) A table browser is provided to search for germplasm by pedigree, origin and subpopulation information.

### The Evolutionary module

Although maize is a well-domesticated and cultivated crop, it still have much to “learn” from their wild relatives, such as the ability to withstand biotic and abiotic stresses (55). In order to provide a general guide for adding new alleles from teosinte into maize breeding programs, ZEAMAP provides selection signals and genetic affinities between maize and its teosinte relatives. The evolutionary selection signals can be browsed graphically through an interactive “Selective signals browser”, which is similar to the aforementioned GWAS viewer but with an additional Y-axis indicating the genetic variance by F_ST_ values (Figure S11A). Signals can also be analyzed in a tabular format (Figure S11B), or viewed in the WashU Epigenome Browser (Figure S11C).

### Epigenetics module

Eukaryotic gene expression has been shown to be altered by three-dimensional DNA interactions, which are affected by chromatin accessibility. Additionally, the modifications of epigenetic states on histones and nucleotides add another layer of control to gene expression regulation (56). These regulatory factors are crucial for the ability of sessile plants to respond to diverse environmental challenges (57). In ZEAMAP, we have collected the chromatin interaction maps associated with RNA polymerase II occupancy and the histone mark H3K4me3 according to the B73 reference genome (6). Open chromatin regions are based on micrococcal nuclease (MNase) digestion (7), histone acetylation and methylation regions, and populational DNA methylation information generated from the third leaves at V3 of the 263 AMP inbred lines (26). This information can be accessed through a tabular data browser or visualized through the WashU Epigenome Browser (Figure 5A). For DNA methylation information from the AMP, customized interfaces were developed to easily select multiple samples with differentially methylated regions (DMRs) in the table browser (Figure 5B) and visualize both DMR and DNA methylation sites in the WashU Epigenome Browser (Figure 5C).

**Figure 5.**
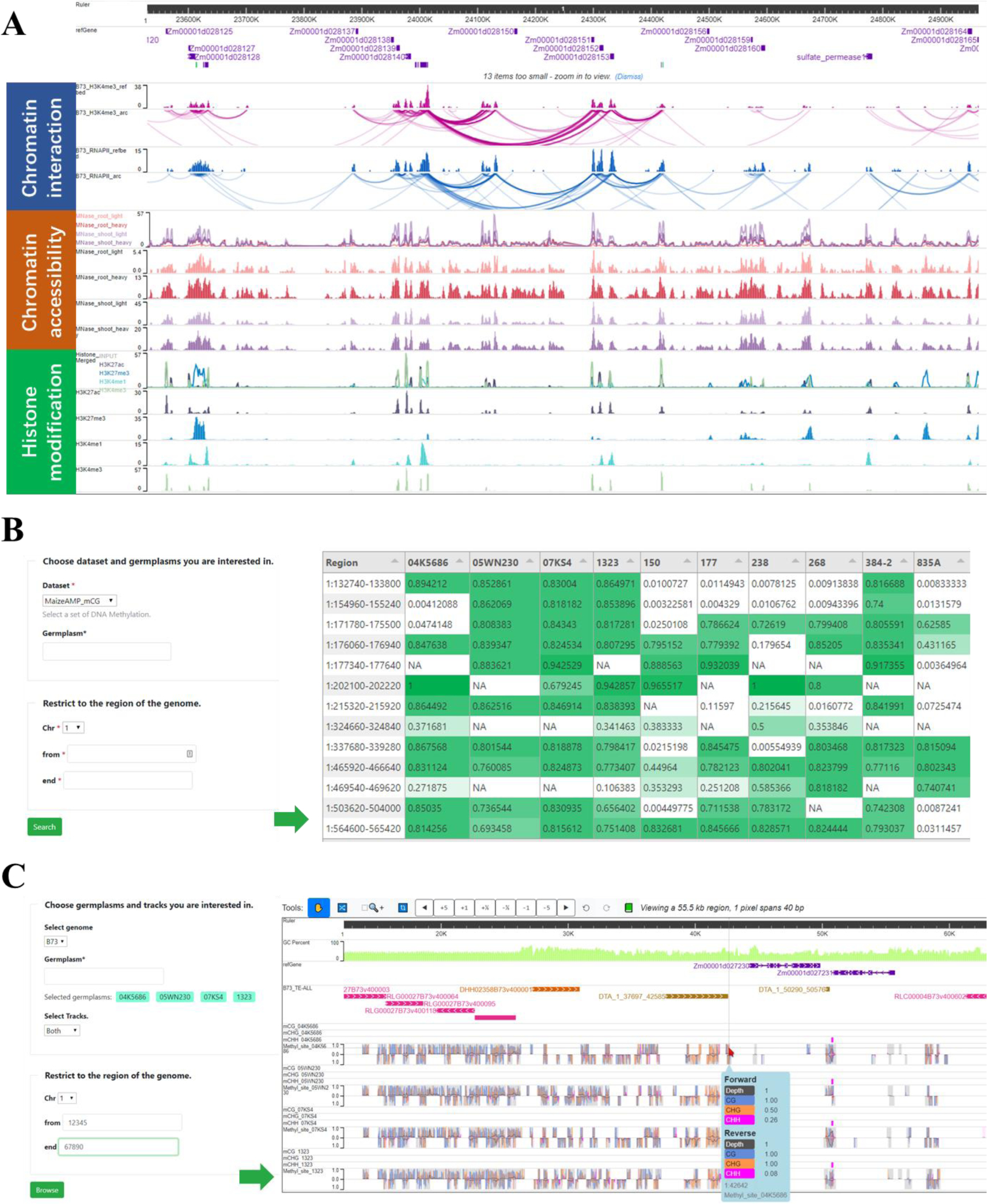
Features of the ZEAMAP Epigenetics module. (A) Schematic of chromatin interaction, chromatin accessibility and histone modification tracks displayed in the WashU Epigenome Browser. (B) Populational DNA methylation table browser. This tool filters population DNA methylation information by the DNA methylation type, germplasm and genomic region of interest, with the resulting matrix displaying DMRs for each selected germplasm within the query region. (C) Interface of the population DNA methylation genome browser. This interface provides options to display DNA methylation information by DMRs or DNA methylation sites of the selected germplasms within specified regions.

### Additional tools

In addition to the aforementioned major biological modules, ZEAMAP also offers several additional tools. The currently available additional tools include a site-wide search engine, BLAST server, a CRISPR browser and an FTP data downloader.

Although there are already independent search tools for several of these analyses, a site-wide search engine powered by Chado is still useful since it enables users to quickly search for all items related to their queries. The ZEAMAP site-wide search engine was built using the Tripal Elasticsearch module (https://github.com/tripal/tripal_elasticsearch), with a search box which is accessible in both the home page and the status bar of each page. The search engine supports advanced search behaviors including wildcards, fuzzy searches, regular expressions and Boolean operators. The search results are also categorized by their entity type in the database (Figure 6A).

**Figure 6.**
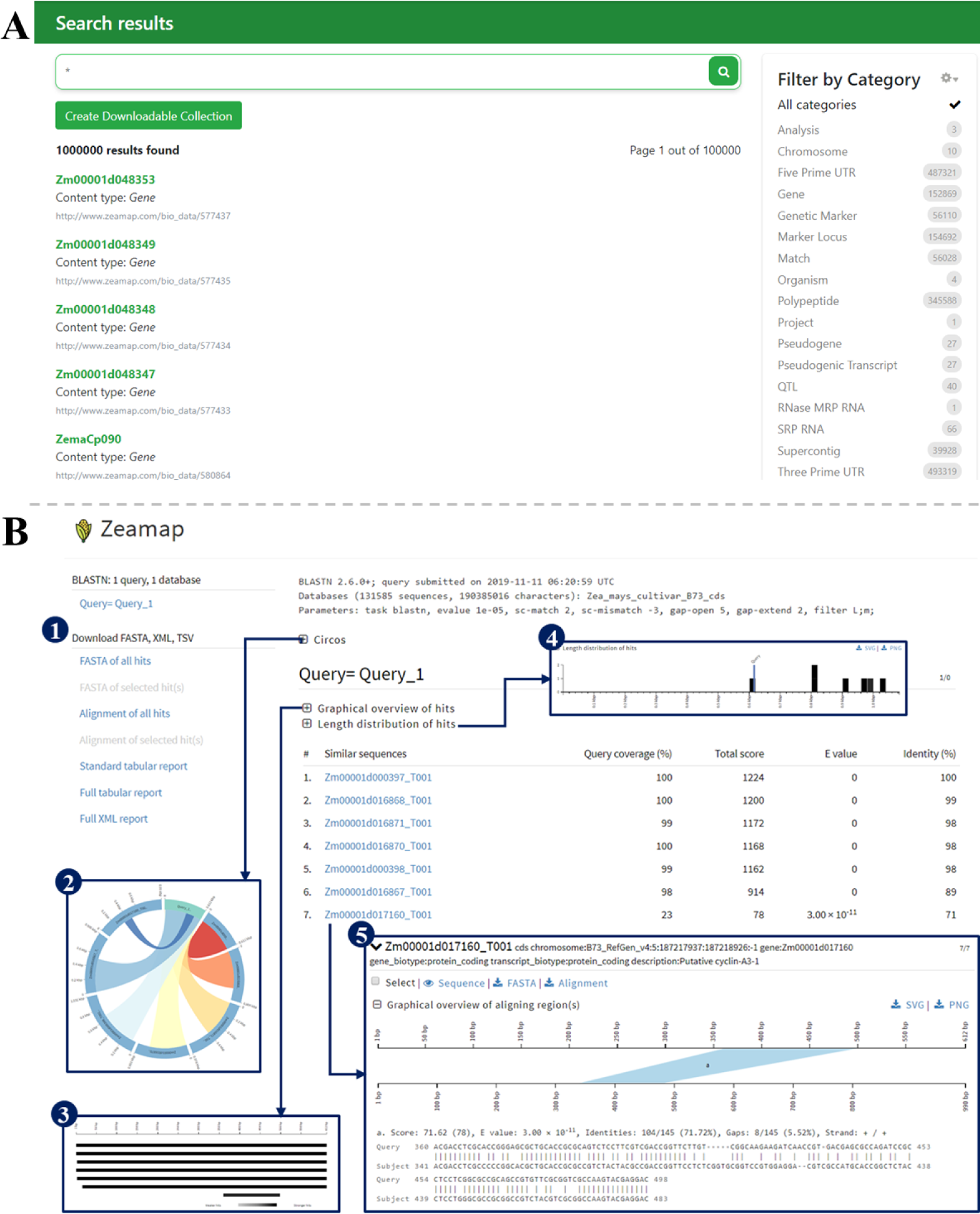
Features of the additional functional tools in ZEAMAP. (A) An example search result by the site-wide search engine queried with an asterisk wildcard. The resulting items are categorized by their feature types and have links to their detailed information pages. (B) An example result of the BLAST server in ZEAMAP. The result page provides download links of sequences① with reports available in different formats. Also included are interactive plot views of each alignment including Circos plots②, NCBI BLAST-like alignment hits visualization③ and length distribution of hits④. Each alignment hit has detailed alignment information, including a graphic view of the aligned regions and detailed alignments⑤.

We have also implemented an instance of NCBI’s BLAST tool in ZEAMAP using SequenceServer (58), which provides a user-friendly interface with text-based and interactive visual outputs (Figure 6B). ZEAMAP currently has BLAST databases for whole genome sequences, mRNAs, CDSs and predicted proteins for each reference genome assembly.

ZEAMAP also includes an imbedded tool to search for reliable single-guide RNAs (sgRNAs) targeting the genes in each maize genome assembly in the database in order to support genome editing experiments using the CRISPR/Cas9 system. The sgRNA information can be browsed tabularly when querying a gene ID or a genomic region, and graphically through JBrowse. Both the tabular and the graphical results provide information about the editing positions and possible off-target genes (Figure S12).

Additionally, we have provided an FTP server to store a backup of all the publicly released datasets used in ZEAMAP through h5ai (https://larsjung.de/h5ai/), an open source file indexer with an enhanced user interface, text preview and directory download.

## Conclusion and future directions

We have created ZEAMAP, a database for maize research and breeding that collects multi-dimensional omics information, including genome assemblies, comparative genomics, transcriptomes, open chromatin, chromatin interactions, genetic variants, phenotypes, metabolomics, genetic maps, genetic mapping loci, population structures and pedigrees, and evolutionary selective signals between teosinte and maize. Most of the datasets were generated from the same maize population, which makes it possible to cross-reference these multi-omics data to support maize research in a more uniform and comprehensive manner. To make the acquisition and analysis of information more effective and flexible, ZEAMAP provides several convenient modules, including a site-wide search function, dataset-specific search tools, a BLAST server, a gene expression pattern analyzer, tabular browsers, genome browsers and specialized visualizers for different datasets.

ZEAMAP will be carefully maintained and continuously updated with the latest genomic and genetic advances. Currently, ZEAMAP has mainly focused on collection, query and visualization of pre-analyzed datasets, with several lightweight online analysis tools embedded. More online analysis tools (software for LD and PCA analyses, for example) will be embedded in ZEAMAP in the near future. Ultimately, we plan to systematically integrate all available omics data and make ZEAMAP a platform to analyze relationships between genotypes and phenotypes in order to predict complex traits for maize researchers and breeders.

## ACKNOWLEDGEMENTS

The authors would like to thank Dr Qing Li and Ms Jing Xu (National Key Laboratory of Crop Genetic Improvement, Huazhong Agricultural University) for providing the DNA methylation data of AMP.

## FUNDING

This research was supported by the National Key Research and Development Program of China (2016YFD0101001, 2016YFD0100303) and the National Natural Science Foundation of China (31525017, 31900494).

