## Supplementary material for "ZEAMAP, a comprehensive database adapted to the maize multi-omics era": Suplemental Figure S1-S12

### Supplementary Files.

Figure S1. Sketch of ZEAMAP database structure.
Figure S2. Species detailed page of *Zea mays* cultivar B73.
Figure S3. Feature search and gene search functions in ZEAMAP.
Figure S4. Features of ZEAMAP variations module.
Figure S5. Search trait function in ZEAMAP.
Figure S6. GWAS table browser in ZEAMAP.
Figure S7. GWAS-Multi-Trait visualization tool in ZEAMAP.
Figure S8. GWAS-Locus visualization tool in ZEAMAP.
Figure S9. eQTL table browser in ZEAMAP.
Figure S10. Genetic marker search functions in ZEAMAP.
Figure S11. Evolution selective signals in ZEAMAP.
Figure S12. Crispr sgRNA function in ZEAMAP.

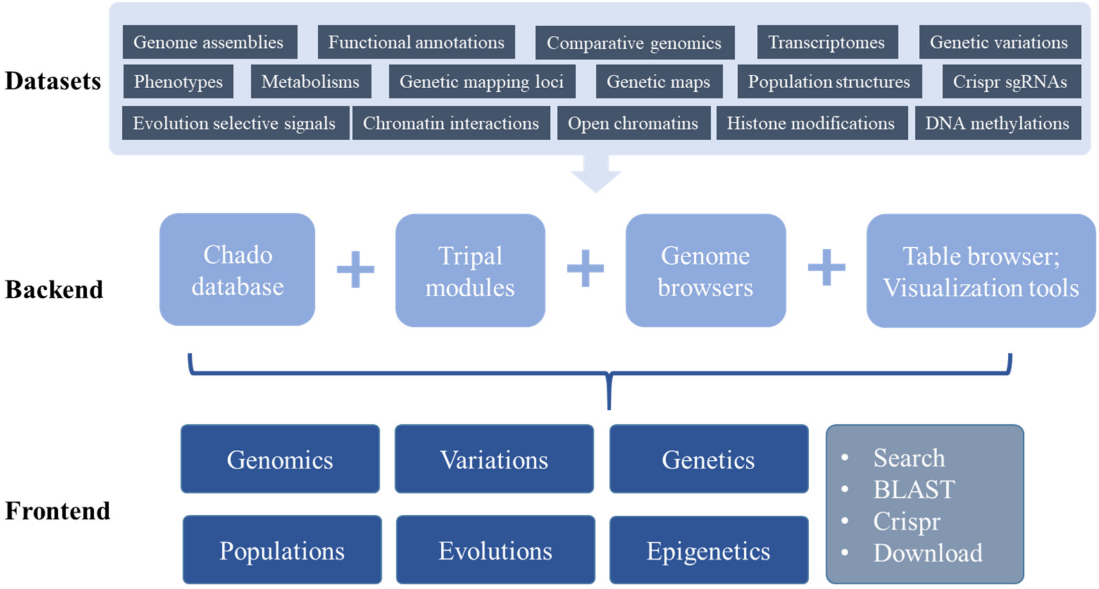


**Figure S1. Sketch of ZEAMAP database structure.** The multi-omics data were stored into database and managed by chado sequence ontology and tripal modules. Two genome browsers, custom table browsers and visualization tools were developed for data querying and visualization. The frontend of ZEAMAP were divided into six main biological sections (Genomics, Variations, Genetics Populations, Evolutions and Epigenetics), and miscellaneous tools including site-wide search engine, BLAST server, Crispr sgRNA searcher and FTP Download server.

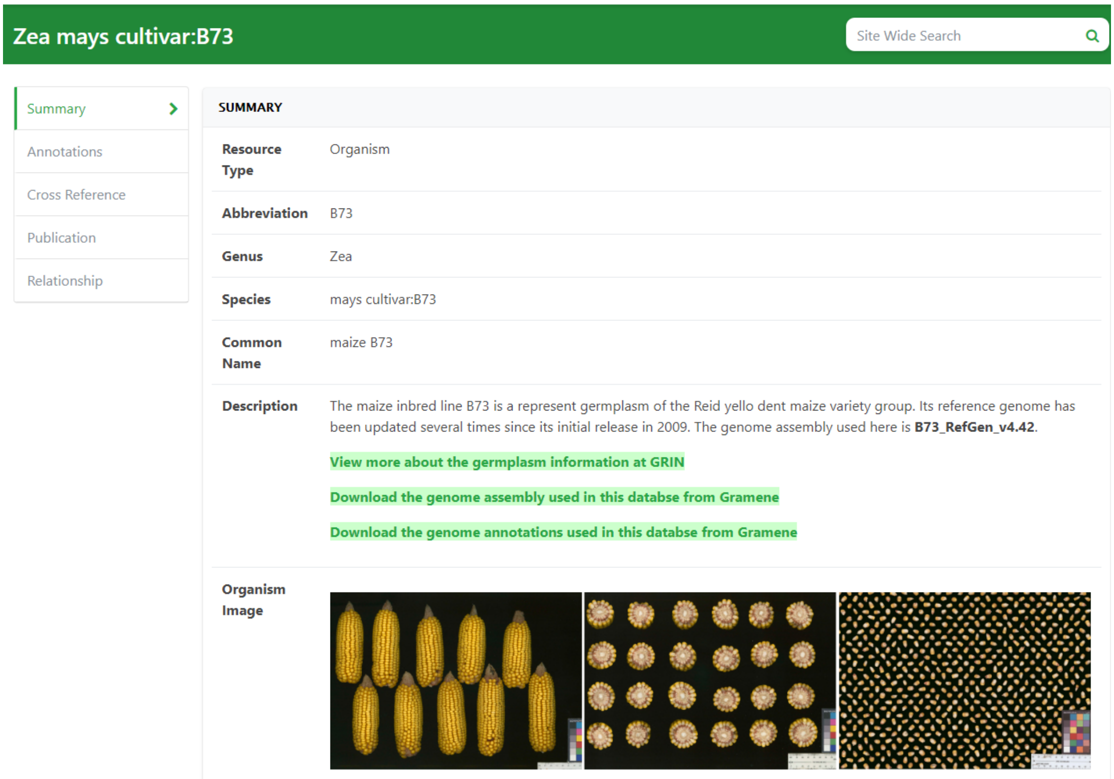


**Figure S2. Species detailed page of *Zea mays* cultivar B73.** The species page shows the general information about the current species/germplasm, including brief introductions, external links to the germplasm information and the accessions of related omics data.

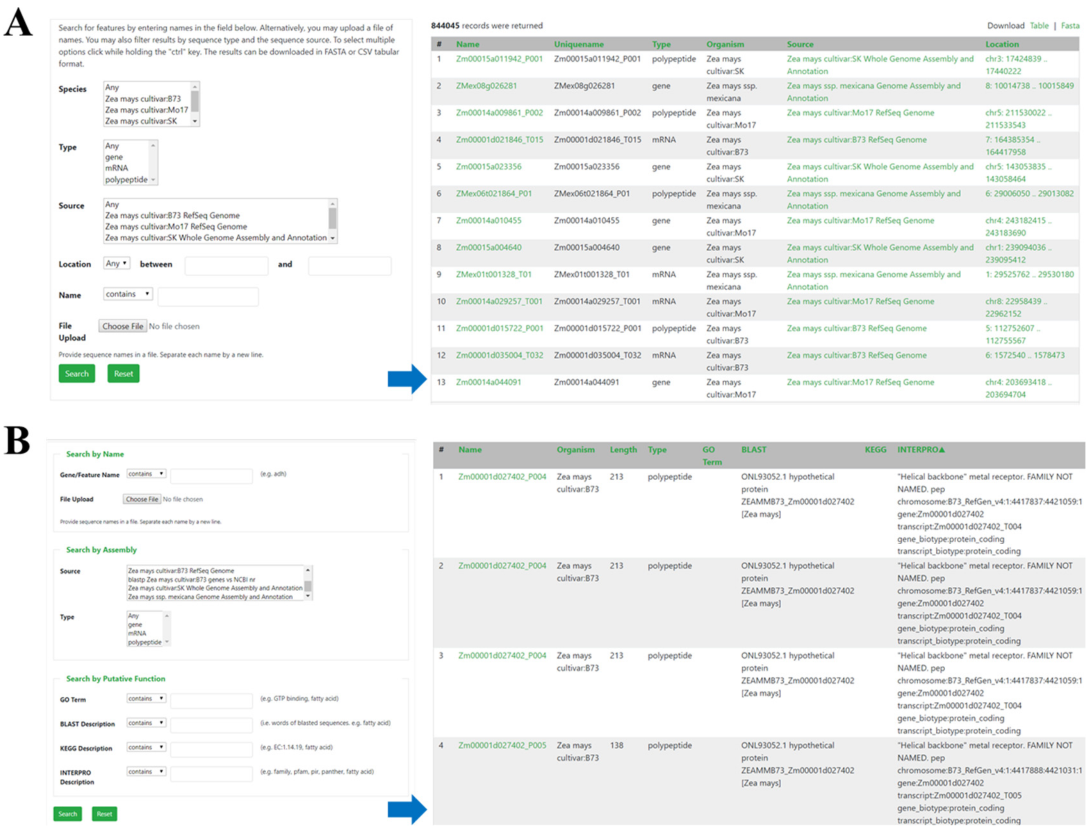


**Figure S3. Feature search and gene search functions in ZEAMAP.** The feature search function (A) in ZEAMAP enables searching for all annotated features on certain genome assemblies by their locations and/or names. The locations of the search results were displayed and linked to genome browser for visualization. The gene search function (B) is dedicated to search for annotated genes, mRNAs and polypeptides by their names and/or their functional annotations, with the detailed functional annotations displayed in the search results for each record.

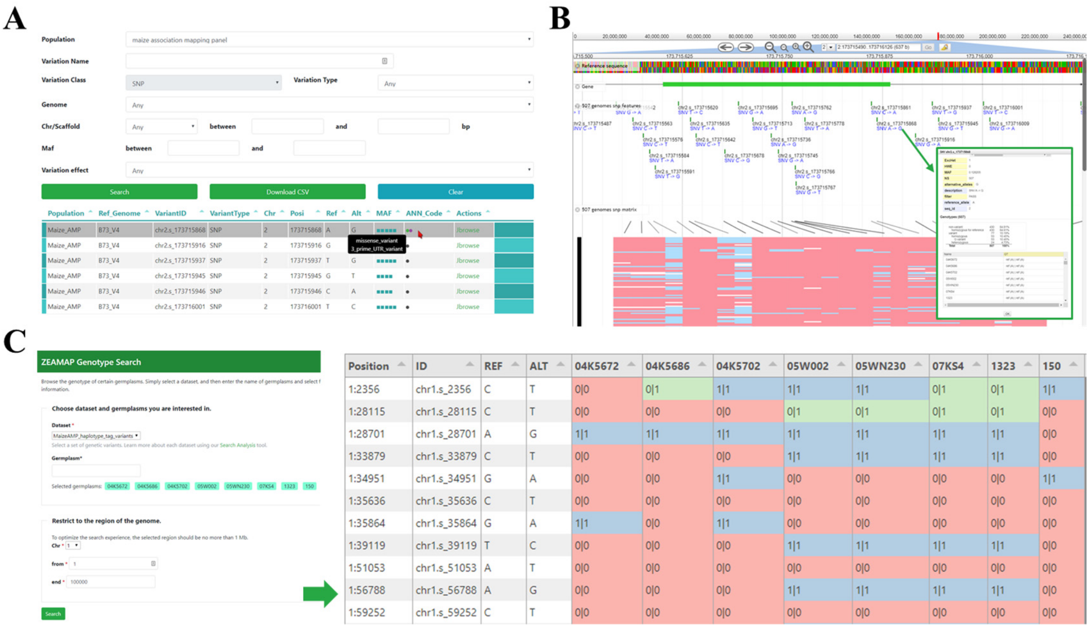


**Figure S4. Features of ZEAMAP variations module.** In ZEAMAP, variants could be displayed tabularly and filtered by IDs, locations and annotated effects (A), with each resulted record has links to its location on Jbrowse (B). The popup for each variant in Jbrowse shows the detailed information including annotations and genotype in each sample (inset in B). Jbrowse has also provided an additional track shows an overview of genotype matrix (red: reference genotype; light blue: homozygous genotype; light pink: heterozygous; grey: no call). (C) ZEAMAP has also provided a "Genotype Search" function to get the genotype information by genomics regions and germplasms of interest, with a resulted matrix shows the variant positions, their IDs, reference and alternative alleles and genotypes in each samples ( "0" for reference allele and "1" for alternative allele; red: homozygous reference genotype; blue: homozygous alternative genotype; green: heterozygous genotype ).

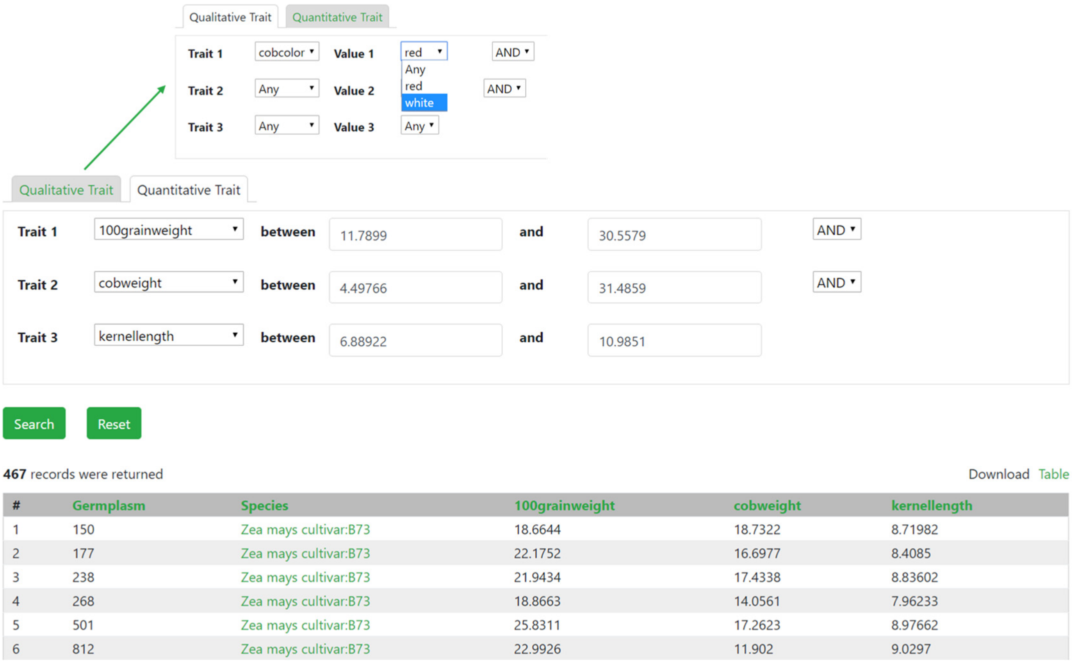

**Figure S5. Search trait function in ZEAMAP.** Both qualitative and quantitative trait could be searched by their trait values with multiple filter conditions supported. The search result shows all the germplasms that passed the filter conditions and their triat values.

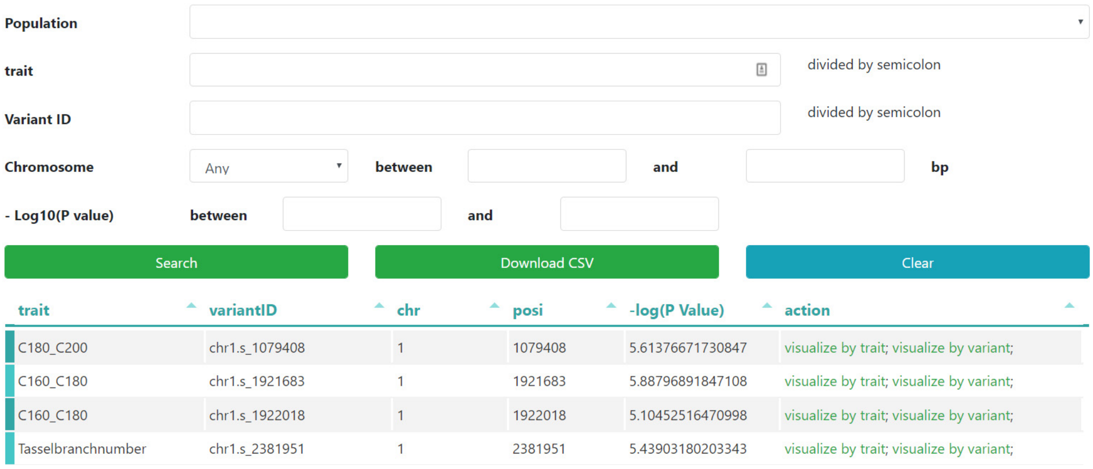


**Figure S6. GWAS table browser in ZEAMAP.** The GWAS signals could be searched by traits, variant IDs and variant locations, and filtered by significant P values. Each record in the search result has links to the GWAS visualization tools.

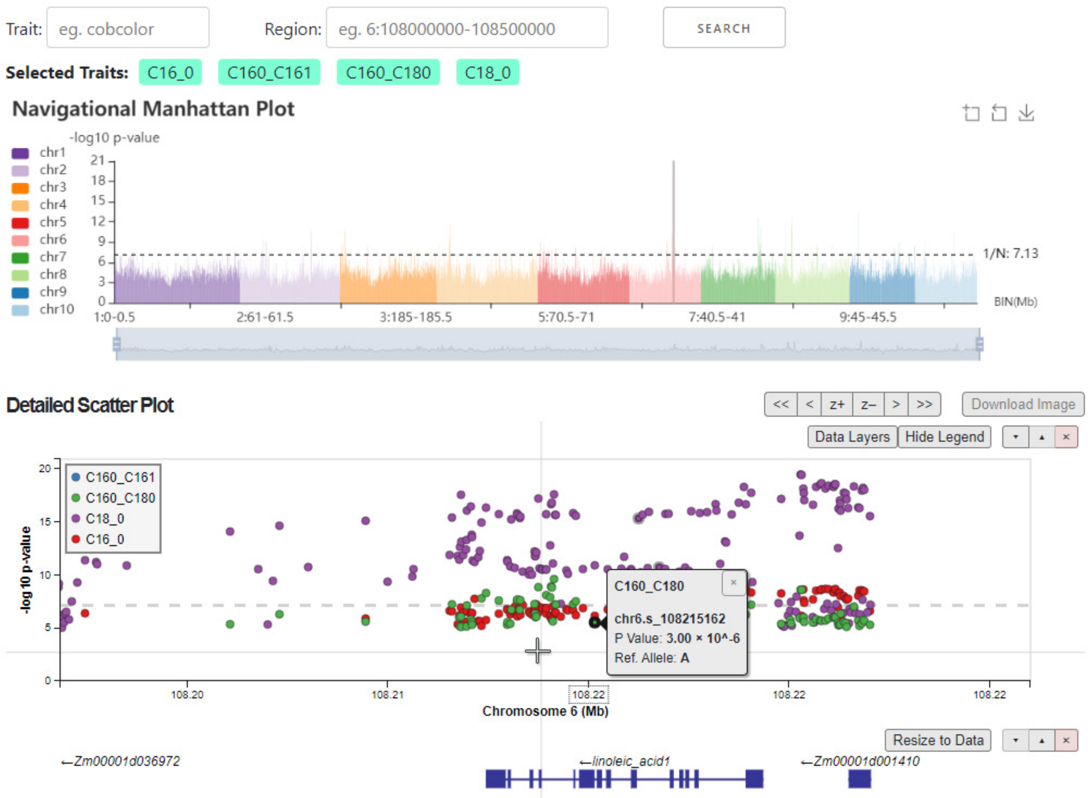


**Figure S7. GWAS-Multi-Trait visualization tool in ZEAMAP.** This tools displays GWAS signals of multiple traits, with logic similar to GWAS-Single-Trait tool (as indicated in Figure 2A ). The only differences are that the traits here are multi-selectable, and the colors in the detailed scatter plot indicate variants for different traits rather than LDs. A “data layers” button has been added in the control panel of the detailed scatter plot to fade, hide, order or remove certain trait layers.

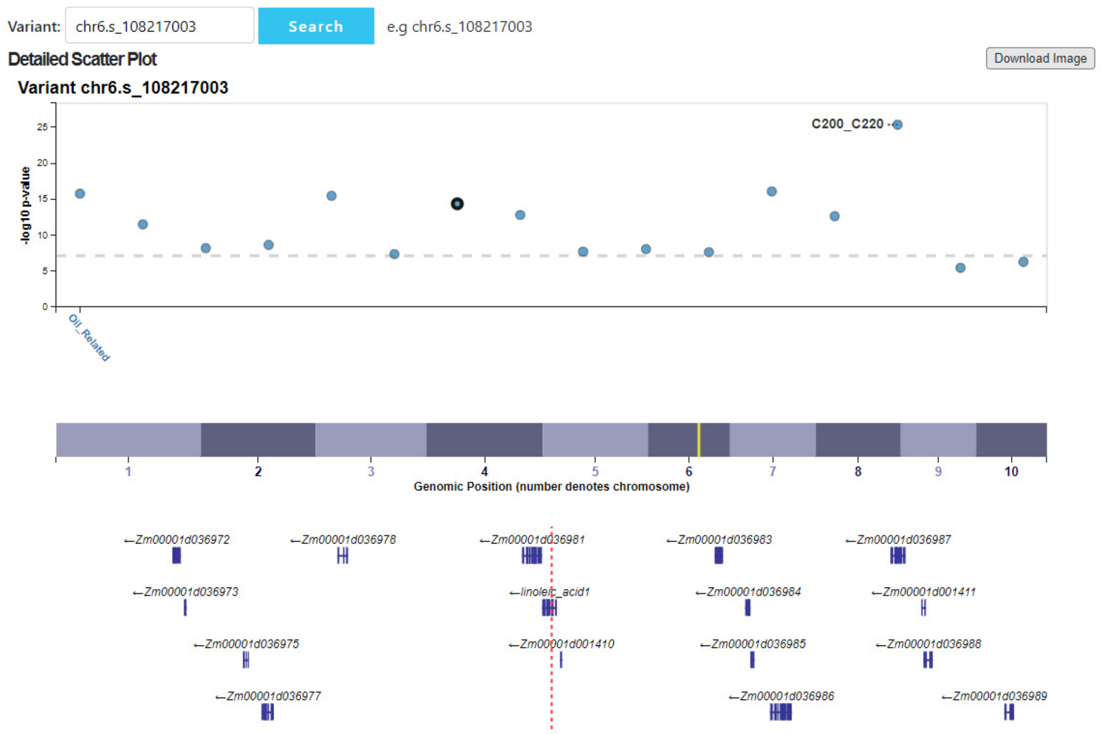


**Figure S8. GWAS-Locus visualization tool in ZEAMAP.** This tool displays all significantly associated signals between the query variant and all available traits, with a yellow highlight line and a red dashed line respectively indicated the general and detailed position of the query variant.

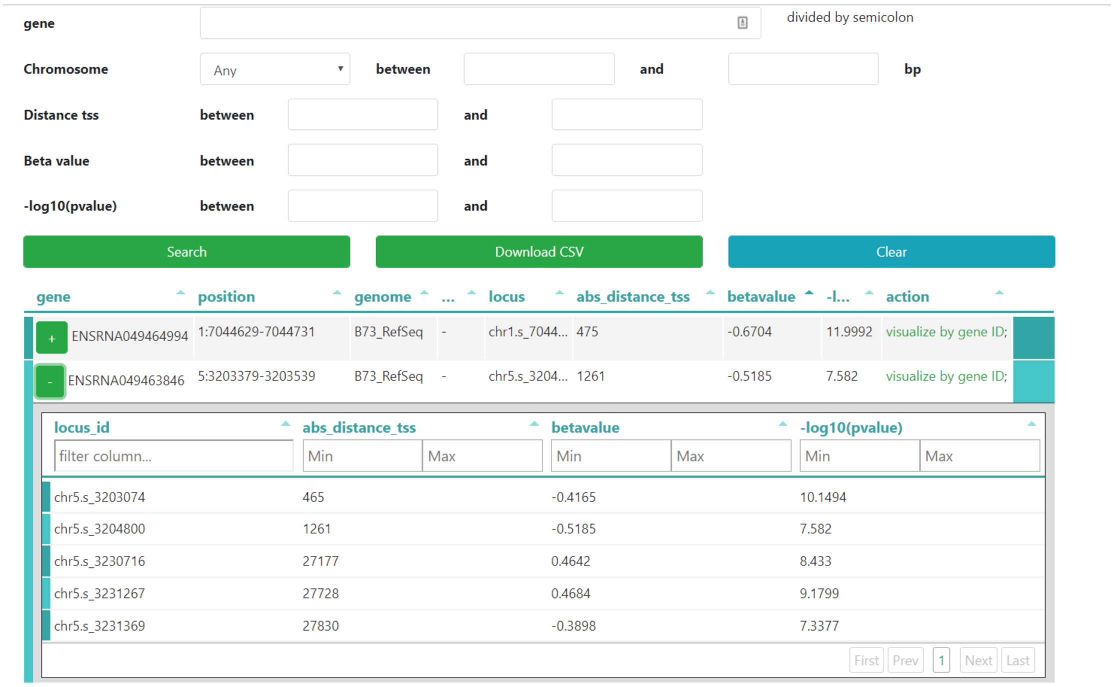


**Figure S9. eQTL table browser in ZEAMAP.** Using this tool, eQTL signals could be filtered by gene IDs and locations, as well as the distance from transcription start site, the effect size (beta value) and the significance (p value) of the most significant variant within each gene. The search result shows one gene per record, with links to the visualization of each gene. Each record has a sub-table which lists all the *cis*-eQTL signals significantly associated with this gene.

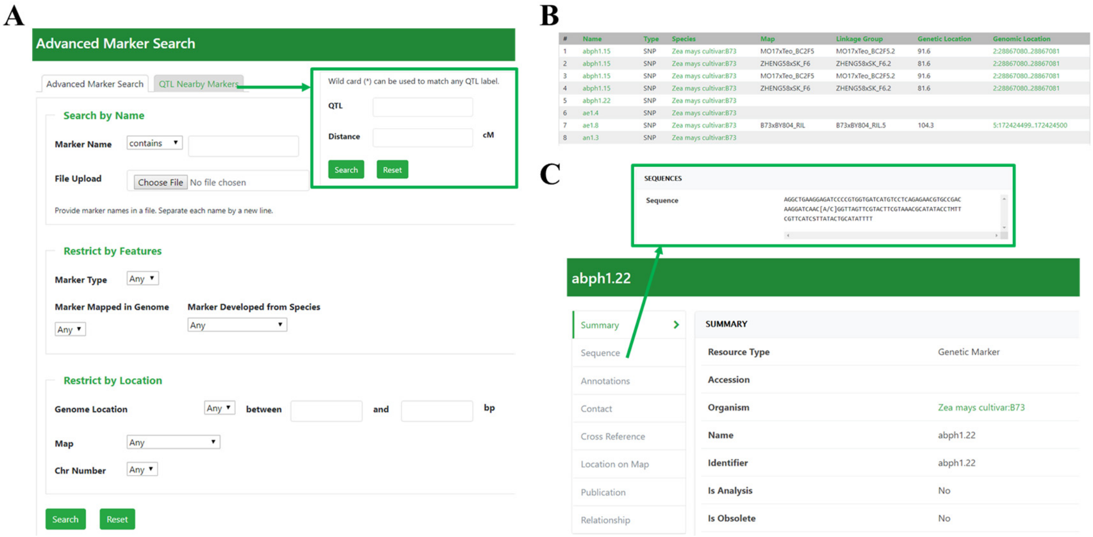

**Figure S10. Genetic marker search functions in ZEAMAP.** The genetic markers in ZEAMAP could be searched by their names, features and locations through "Advanced Marker Search" function (A) or by their distances with certain QTL through "QTL Nearby Marker" function (inset in A). The search result lists general information of genetic and physical locations for each record, with physical locations linked to Jbrowse visualization (B). Click on each marker name would lead to the detailed page for this marker including the flanking sequence and the location on maps (C).

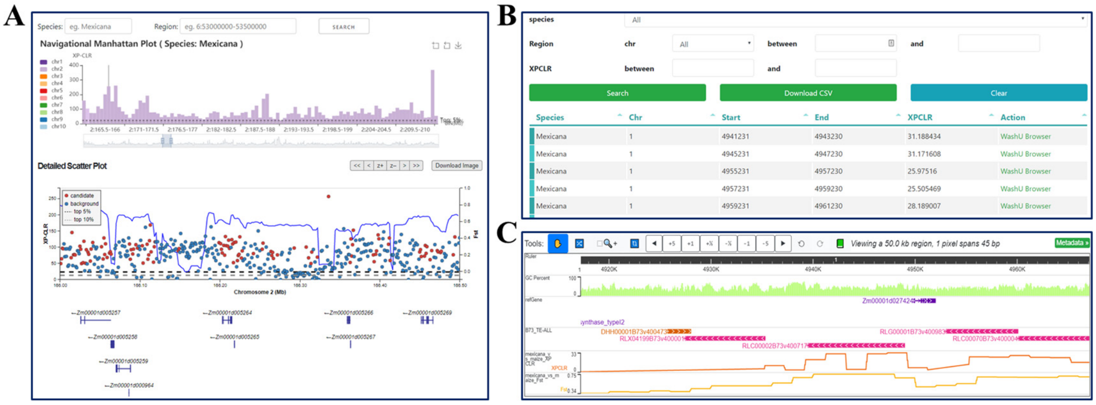


**Figure S11. Evolution selective signals in ZEAMAP.** (A) Selective signals browser. The interactive histogram shows the distribution of XPCLR values within 500Kb windows along the chromosomes, with Y-axis indicated the maximum XPCLR scores within each window. The detailed scatter plot shows the detailed selective signals (dots) and the $F_{st}$ values (blue line) within the selected region, with two horizontal dash lines indicated the top 5% and top 10% XPCLR value cutoffs. (B) A table browser was provided to search for selective signals by teosinte sub-species and genomic regions, each resulted record has links to its position on (C) WashU Epigenome Browser.

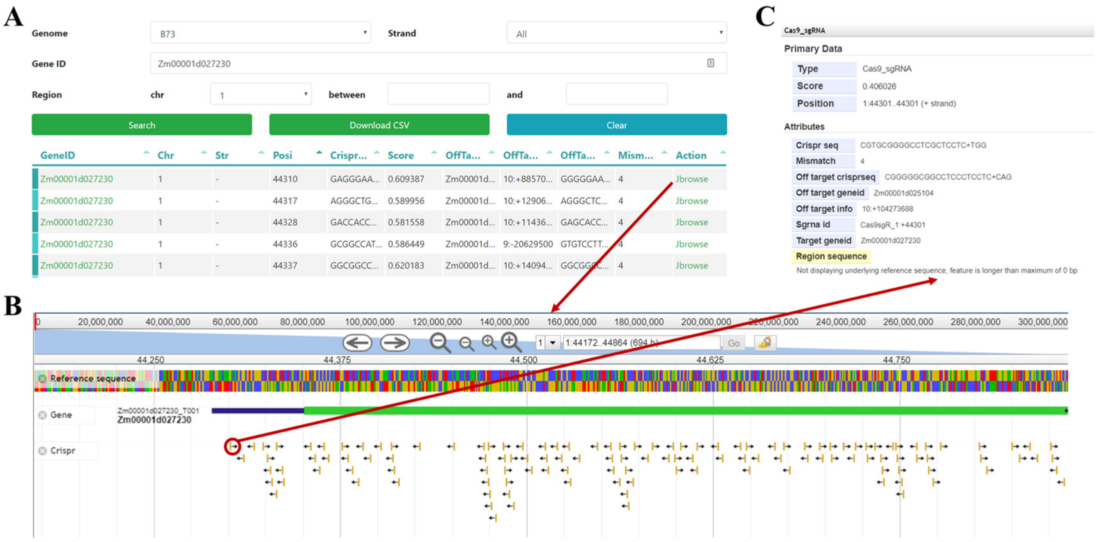

**Figure S12. Crispr sgRNA function in ZEAMAP.** The sgRNA information could be searched by IDs and genome locations of target genes through table browser (A), the resulted records have links to their locations on Jbrowse (B) and the detail page of each Cas9_sgRNA element on Jbrowse shows the information of target and off-target genes (C).
